# Tumour specimen cold ischemia time impacts molecular cancer drug target discovery

**DOI:** 10.1101/2024.05.23.595517

**Authors:** Silvia von der Heyde, Nithya Raman, Nina Gabelia, Xavier Matias-Guiu, Takayuki Yoshino, Yuichiro Tsukada, Gerry Melino, John L. Marshall, Anton Wellstein, Hartmut Juhl, Jobst Landgrebe

## Abstract

Tumour tissue collections are used to uncover pathways associated with disease outcomes that can also serve as targets for cancer treatment, ideally by comparing the molecular properties of cancer tissues to matching normal tissues. The quality of such collections determines the value of the data and information generated from their analyses including expression and modifications of nucleic acids and proteins. These biomolecules are dysregulated upon ischemia and decomposed once the living cells start to decay into inanimate matter. Therefore, ischemia time before final tissue preservation is the most important determinant of the quality of a tissue collection. Here we show the impact of ischemia time on tumour and matching adjacent normal tissue samples for mRNAs in 1,664, proteins in 1,818 and phosphoproteins in 1,800 cases (tumour and matching normal samples) of four solid tumour types (CRC, HCC, LUAD and LUSC NSCLC subtypes). In CRC, ischemia times exceeding 15 minutes impacted 12.5% (mRNA), 25% (protein) and 50% (phosphosites) of differentially expressed molecules in tumour versus normal tissues. This hypoxia- and decay-induced dysregulation increased with longer ischemia times and was observed across tumour types. Interestingly, the proteomics analysis revealed that specimen ischemia time above 15 minutes is mostly associated with a dysregulation of proteins in the immune response pathway and less so with metabolic processes. We conclude that ischemia time is a crucial quality parameter for tissue collections used for target discovery and validation in prognostic cancer research.

## 1 Introduction

The identification rate of novel protein targets in the context of solid cancer therapy has decreased over the last ten years.

One of the major concepts in identifying functional proteins as oncogenic targets is the detection of alterations that lead to aberrant signalling in cancer-relevant pathways. A core principle of today’s onco-pharmacological targeting approach is detecting and applying therapeutics that antagonize oncogenic protein function and thereby significantly improve overall survival [1]. An important strategy to systematically identify new target proteins for cancer treatment is to build a cancer registry combining tissue and meta-data collection, a strategy pursued, for example, by the TCGA consortium.^1^ To this end, cancer and ideally also matching normal adjacent tissue are collected and analysed to compare the molecular characteristics of the two tissue types. The two parameters which have the strongest impact on the quality of tissue registries are the asservation method and the so-called cold ischemia time (in short ‘ischemia time’), which denotes the time it takes to freeze the tissue for permanent storage after its removal from the body during surgery.

There is an ongoing debate about the consequences of ischemia on the quality of biomolecules, especially in the context of characterising the molecular entirety of tissues or cells (so-called ‘omics’). The corresponding findings vary widely [2, 3, 4] due to insufficient numbers of samples which leads to an insufficient power to detect the impact of ischemia time on the molecular composition of the materials. Hence, the impact of ischemia time on the molecular characteristics of the tissue under investigation remains unclear. Unlike any previous study, we use adequate sample sizes and statistical methods to investigate the impact of ischemia time on target discovery.

Here, we focus on the impact of ischemia time on differential expression of mRNAs, proteins and phosphoproteins using a carefully curated collection of patient-derived fresh-frozen tissue samples and related multiomic data. This data base contains the breadth of molecular characteristics of tumour and normal adjacent tissues from multiple cancer types. Specimens were analysed at the genomic, transcriptomic (TRX), proteomic (PTX) and phosphoproteomic (PPX) level with focus on colon cancer (CRC). Analyses were expanded to hepatocellular carcinoma (HCC) and non-small cell lung epithelial cancer (NSCLC), covering lung adenocarcinoma (LUAD) and lung squamous cell carcinoma (LUSC).

To generate a baseline in our studies we set an initial filter to capture biomolecules differentially expressed in the group of samples with shortest ischemia time available (*t* < 10 min.). We then analyse the changes in expression of these biomolecules over time with an emphasis on the impact of ischemia time on the target identification process, rather than trying to model the tissue decay under ischemia.

We found that DNA characteristics are to a large extent unaffected even by the longest ischemia times of samples collected. In contrast, the relationship of tumour and normal tissue mRNA, protein and phosphoprotein expression is affected at increasing severity (mRNA < protein < phosphoproteins). Based on the analyses we propose an ischemia time cut-off threshold at 12 minutes that enables a sufficient amount of tissues to be collected while avoiding a dilution of signals for biomolecules of interest.

## 2 Results

### 2.1 Cancer samples and differential expression

Data sets and samples used in the analysis are shown in Table 1, tumour stages are shown in Table 2. Details on the samples can be found in the methods section.

**Table 1:**
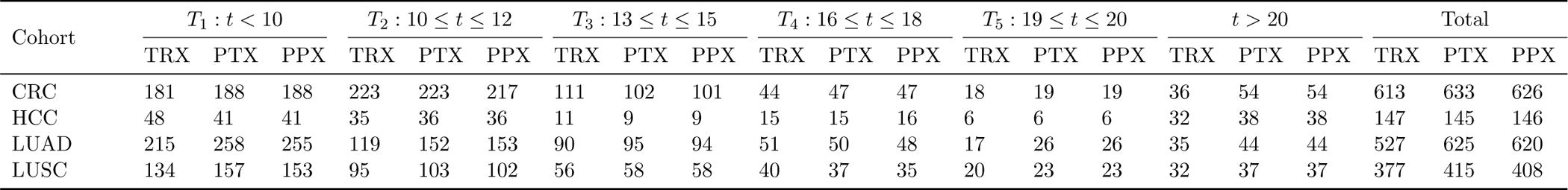
Number of tumour samples per ischemia time interval (min.) with specific omic data.

| Cohort | $T_1 : t < 10$ | | | $T_2 : 10 \leq t \leq 12$ | | | $T_3 : 13 \leq t \leq 15$ | | | $T_4 : 16 \leq t \leq 18$ | | | $T_5 : 19 \leq t \leq 20$ | | | $t > 20$ | | | Total | | |
| --- | --- | --- | --- | --- | --- | --- | --- | --- | --- | --- | --- | --- | --- | --- | --- | --- | --- | --- | --- | --- | --- |
|  | TRX | PTX | PPX | TRX | PTX | PPX | TRX | PTX | PPX | TRX | PTX | PPX | TRX | PTX | PPX | TRX | PTX | PPX | TRX | PTX | PPX |
| CRC | 181 | 188 | 188 | 223 | 223 | 217 | 111 | 102 | 101 | 44 | 47 | 47 | 18 | 19 | 19 | 36 | 54 | 54 | 613 | 633 | 626 |
| HCC | 48 | 41 | 41 | 35 | 36 | 36 | 11 | 9 | 9 | 15 | 15 | 16 | 6 | 6 | 6 | 32 | 38 | 38 | 147 | 145 | 146 |
| LUAD | 215 | 258 | 255 | 119 | 152 | 153 | 90 | 95 | 94 | 51 | 50 | 48 | 17 | 26 | 26 | 35 | 44 | 44 | 527 | 625 | 620 |
| LUSC | 134 | 157 | 153 | 95 | 103 | 102 | 56 | 58 | 58 | 40 | 37 | 35 | 20 | 23 | 23 | 32 | 37 | 37 | 377 | 415 | 408 |

**Table 2:** Number of tumour samples per cancer stage group with specific omic data.

| Cohort | I |  |  | II |  |  | III |  |  | IV |  |  | NA |  |  | Total |  |  |
| --- | --- | --- | --- | --- | --- | --- | --- | --- | --- | --- | --- | --- | --- | --- | --- | --- | --- | --- |
|  | TRX | PTX | PPX | TRX | PTX | PPX | TRX | PTX | PPX | TRX | PTX | PPX | TRX | PTX | PPX | TRX | PTX | PPX |
| CRC | 65 | 60 | 60 | 229 | 251 | 248 | 195 | 201 | 199 | 121 | 118 | 116 | 3 | 3 | 3 | 613 | 633 | 626 |
| HCC | 31 | 33 | 34 | 18 | 22 | 22 | 18 | 17 | 17 | 7 | 7 | 7 | 73 | 66 | 66 | 147 | 145 | 146 |
| LUAD | 232 | 293 | 291 | 129 | 145 | 144 | 133 | 143 | 141 | 24 | 33 | 33 | 9 | 11 | 11 | 527 | 625 | 620 |
| LUSC | 126 | 151 | 147 | 125 | 130 | 130 | 116 | 119 | 116 | 6 | 8 | 8 | 4 | 7 | 7 | 377 | 415 | 408 |

We defined the ischemia time reference group as the group of samples with ischemia times less than 10 minutes, the shortest time possible for the collection and storage of a sufficient number of samples. To obtain differential biomolecule expression for mRNAs, proteins and phoshoproteins for the shortest ischemia time group, we normalised the expression data and selected the differentially expressed biomolecules using non-parametric Wilcoxon tests for paired samples to take into account the non-normal distribution of the data [5] as described in the methods part. We adjusted the resulting P-values for multiple testing and selected the biomolecule sets for mRNA, protein, and phosphoprotein using an *α*_fdr_ level of 0.01 and an effect boundary using the 5th and 95th percentiles of the effect distribution. This yields 1,948 differentially expressed mRNAs, 794 proteins and 1,846 phosphosites on which we focussed in the CRC cohort. Analogously, we inferred 1,870 differentially expressed mRNAs, 523 proteins and 388 phosphosites in the HCC cohort. In the LUAD cohort, we inferred 1,951 differentially expressed mRNAs, 805 proteins and 2,217 phosphosites, and in the LUSC cohort, we inferred 1,950 differentially expressed mRNAs, 798 proteins and 1,919 phosphosites on which we focussed in the subsequent analyses.

### 2.2 Differential expression over time in CRC

We performed a detailed analysis of differential (tumour versus normal adjacent tissue) expression over time in CRC, and then applied the most relevant analyses to the other cancer types listed in Table 1.

To evaluate whether DNA sequences remain unaffected by ischemia times as reported by others [6], we analysed protein sequences for affecting mutations (PAM), the presence or absence of gene deletions or amplification as well as gene truncations by comparing the shortest available ischemia time tissue group with the longer ischemia time groups for each genomic locus (see Methods). At *α*_fdr_ = 0.01, only 0.09% of the proteins showed a signal in the PAM submodality, and no change was found in the other DNA-derived submodalities. This is likely based on random somatic difference in the genomic sequence of the patients of the various time groups (see Discussion).

To obtain an overview of the effect of ischemia time on the selected biomolecules, we initially partitioned the samples into groups of 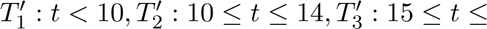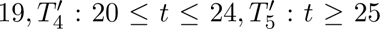 minutes of ischemia time duration intervals. We then used hierarchical clustering to assess the changes in biomolecule expression for the three omic modalities as described in the methods section. Figure 1 shows the results for the phosphoprotein modality for *k* = 10 clusters (the corresponding plots for the other two expression modalities are shown in Supplementary Figure S1).

**Figure 1:**
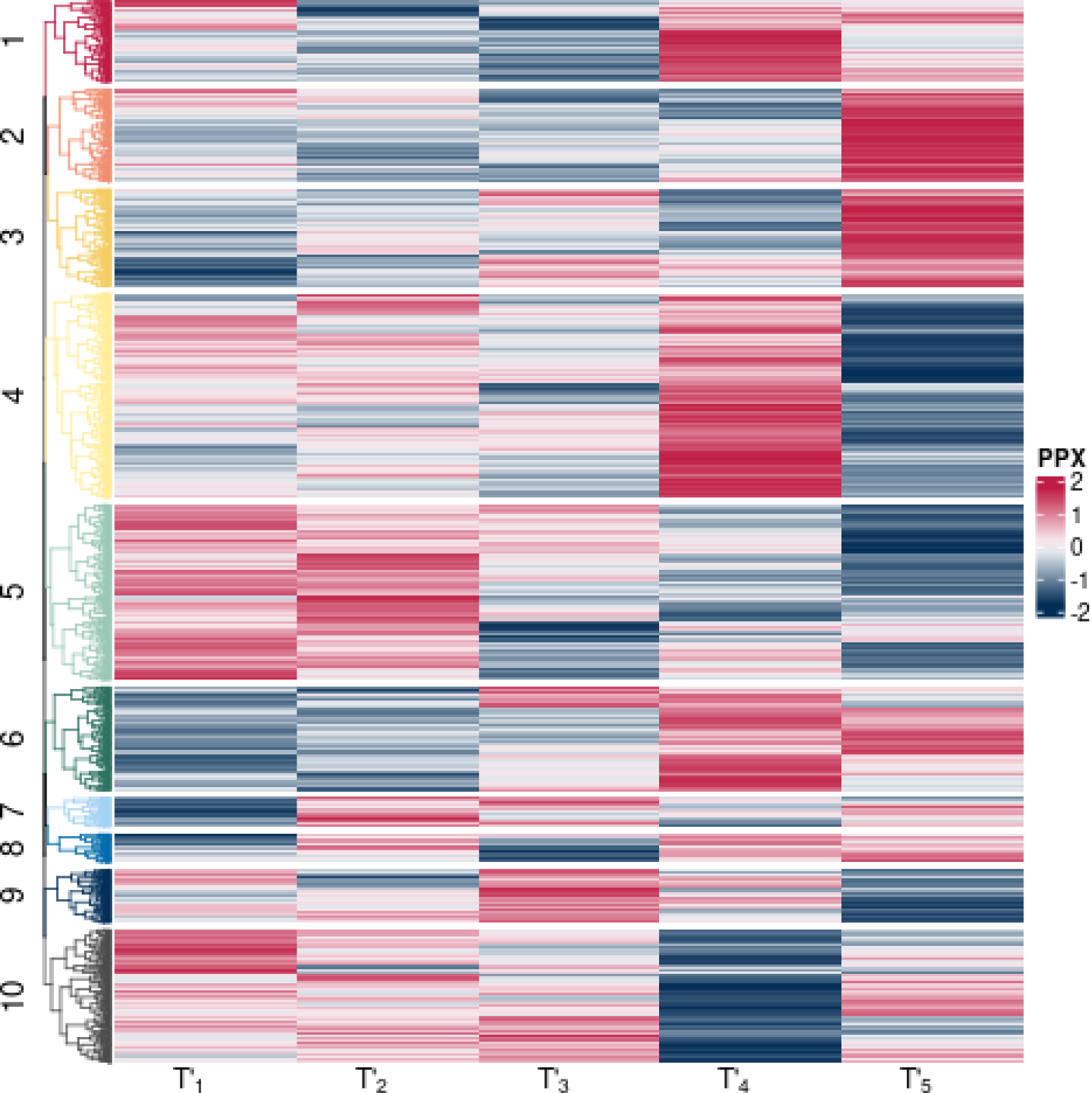
Differential expression of the selected phosphosites in 5-minute intervals (abscissa) grouped into ten clusters (ordinate) in CRC. Colours indicate log_2_-fold mean expression differences (phosphosite-wise standardised) between tumour and normal tissue. Red: upregulation in tumour, blue: downregulation.

The effect magnitude and significance seem to increase for times over 20 minutes. To confirm this impression, we tested for differences in differential expression over time in each modality using a 2 × 5 factorial ANOVA-like design with the tissue types (tumour and normal) as first factor and the time groups 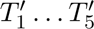 as second factor using the non-parametric Scheirer-Ray-Hare test also assessing interaction effects between the two main effects. Table 3A shows the phosphosites per cluster with significant tissue, time or interaction effects. The three effects on display are the two main effects of tissue and time difference and the interaction effect of the two variables.

**Table 3:** Alterations in phosphosite expression over time across tissue types.

| **A: Long duration ischemia time groups** |  |  |  | **B: Refined ischemia time groups** |  |  |  |
| --- | --- | --- | --- | --- | --- | --- | --- |
| Significant ($\alpha_{\text{fdr}} = 0.05$) effect for | | | | Significant ($\alpha_{\text{fdr}} = 0.05$) effect for | | | |
| Cluster | Tissue | Time | Interaction | Cluster | Tissue | Time | Interaction |
| 1a | 147 | 45 | 0 | 1b | 115 | 16 | 0 |
| 2a | 169 | 63 | 3 | 2b | 293 | 51 | 2 |
| 3a | 177 | 61 | 6 | 3b | 283 | 35 | 2 |
| 4a | 369 | 129 | 2 | 4b | 95 | 11 | 0 |
| 5a | 317 | 101 | 12 | 5b | 136 | 25 | 0 |
| 6a | 188 | 68 | 8 | 6b | 89 | 10 | 0 |
| 7a | 54 | 11 | 2 | 7b | 442 | 80 | 1 |
| 8a | 52 | 16 | 0 | 8b | 189 | 21 | 2 |
| 9a | 97 | 35 | 0 | 9b | 74 | 15 | 0 |
| 10a | 242 | 76 | 6 | 10b | 105 | 20 | 2 |
| Totals | 1812 | 605 | 39 | Totals | 1821 | 284 | 9 |

Table 3 confirms that across all clusters (note that the clusters in both panels do not contain identical phosphosites since the data vectors per phosphosite differ between the panels) of the original time groups (panel A), 33% (605/1814) of the phosphosites display a significant time effect. We therefore repartitioned the available samples into the shortest ischemia group (samples with ischemia time shorter than 10 minutes), and into further groups of 3 minute intervals from *t* ≥ 10 to 20 minutes (*T*_1_ … *T*_5_) as shown in Table 1 in order to quantify which ischemia time point to use as cut-off (we call these *refined* time groups). With these refined groups (Table 3, panel B) which exclude longer ischemia times, the time effect halves to 16% (284/1823) of the phosphosites.^2^ There are also less interaction effects. The trends for mRNA and protein modalities are comparable, though less pronounced (see Supplementary Figure S1 and Supplementary Tables S1 and S2). But even though the effect is least pronounced in mRNA, to perform valid multiomics analyses, we need an ischemia time limit that is the same for all modalities. Thus, we aim for a limit that is optimal for the most sensitive type of molecule (phosphate groups), which must be under 20 minutes.

Before investigating differential biomolecule expression, we analysed the distribution of the underlying expression levels in normal and cancer tissue samples of the groups *T*_1_ … *T*_5_. We specifically focused on biomolecules with extreme expression values which could play an important role in cancer. We regarded biomolecules with expression values outside the (*Q*_2.5_*, Q*_97.5_) interval as extremely expressed. We determined the relative loss of such extreme biomolecules of reference group *T*_1_ in other ischemia time groups. In detail, we counted how many biomolecules with extreme expression values outside (*Q*_2.5_*, Q*_97.5_) of *T*_1_ were not detected outside the (*Q*_2.5_*, Q*_97.5_) intervals of the other groups *T*_2_ … *T*_5_ any more. For example for *T*_1_ and *T*_2_, this relative loss is

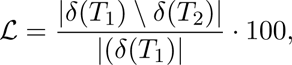

where *δ* computes the biomolecules outside the interval (*Q*_2.5_*, Q*_97.5_) and | · · · | gives the set size. Table 4 shows the loss rates for tumour and normal tissues for the time group average. The vanishing biomolecules fall from outside (*Q*_2.5_*, Q*_97.5_) into the interval in the time groups *T*_2_ … *T*_5_.

**Table 4:** Loss L of biomolecules outside the (*Q*_2.5_*, Q*_97.5_) interval.

| Modality | Tissue | Biomolecule loss [%] vs. $T_1$ | | | |
| --- | --- | --- | --- | --- | --- |
| | | $T_2$ | $T_3$ | $T_4$ | $T_5$ |
| mRNA | Normal | 27 | 32 | 39 | 21 |
|  | Tumour | 23 | 28 | 45 | 27 |
| Protein | Normal | 24 | 25 | 36 | 56 |
|  | Tumour | 20 | 23 | 27 | 42 |
| Phosphoprotein | Normal | 49 | 64 | 72 | 94 |
|  | Tumour | 46 | 60 | 64 | 81 |

The loss of biomolecule sets in group *T*_2_ is striking for all types of analysed biomolecules, but especially for phosphosites. Notably, the loss of the most highly expressed mRNAs is slower than for proteins and even more so than for phosphoproteins, where almost 50% of the most highly expressed sites are lost in a short interval time between 10 and 12 minutes of ischemia. In mRNA, there seems to be a recovery of the initial expression pattern after the longest ischemia duration, an artefact which may be explained by the degradation and altered detection of the molecules. This has to be taken into account when looking at differential biomolecule expression.

#### 2.2.1 Temporal patterns of refined time series

To evaluate the influence of ischemia on differential expression patterns, we next analysed the shorter interval groups *T*_1_ … *T*_5_ using a Dirichlet process Gaussian process mixture model (DPGP, details see Material and methods). This non-parameteric time series analysis technique jointly models data clusters with a Dirichlet process and temporal dependencies with a Gaussian process.^3^

Figure 2 shows the eleven clusters obtained in CRC for the protein modality, which we selected out of the three available expression modalities because it allows a deeper biological interpretation (analogous time series analyses can be generated for each modality and tumour type).

**Figure 2:**
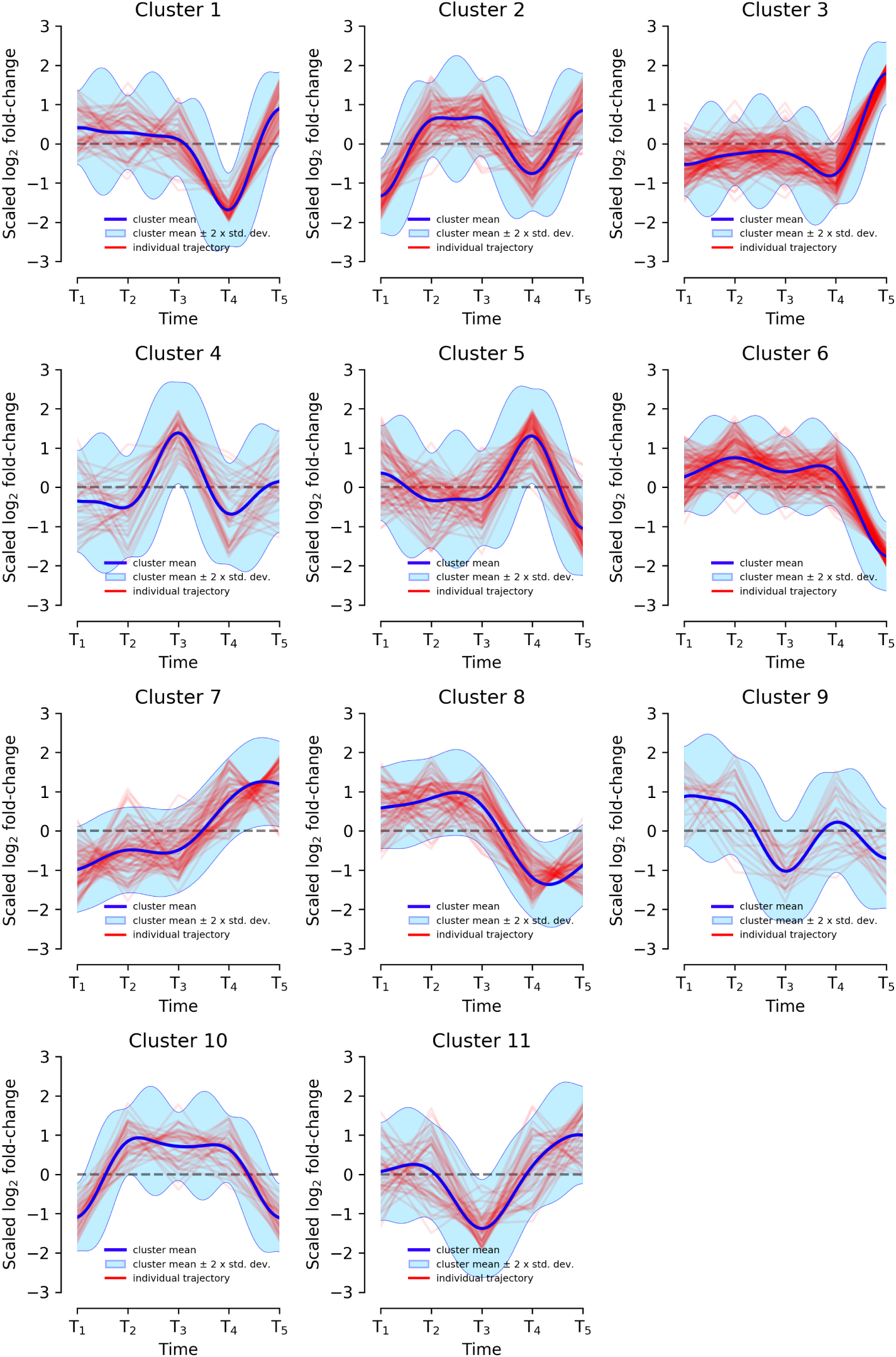
Differential expression of the selected proteins in refined time groups (*T*_1_ … *T*_5_ as 3-minute intervals on the abscissa) grouped into 11 clusters. The ordinates of each cluster show the normalised differential expression. The blue line shows the cluster mean expression. The red lines indicate the individual protein expression levels. The shaded blue area indicates the cluster mean ±2 · *σ*.

We expect various patterns of protein expression variance due to ischemia with proteins losing or gaining expression levels under the effect of the ischemia-induced decay of the tissue.

**Biological cluster interpretation** To interpret the biological meaning of the protein expression patterns, we used a Wilcoxon test to identify proteins with differential expression from one time group to another (*µ_T_*_2_ −*µ_T_*_1_ *, …, µ_Tk_*_+1_ −*µ_Tk_, k* = 2 … 4) excluding effects inside the interval (2^−1/2^, 2^1/2^).

We then mapped the genes related to these differentially expressed proteins to KEGG^4^ and performed a GO enrichment analysis for biological pathways [7]. We could not find cluster-specific relevant biological patterns, maybe because ischemia regulation cannot be revealed by DPGP-clustering on our pseudo-time-series (see footnote 5), but we still found interesting overall patterns. The analysis revealed acute inflammatory response and metabolism as the most significantly up- and downregulated pathways, respectively. Table 5 shows the amount of significantly differentially expressed proteins per pathway. Quite surprisingly, the data show a strong upregulation of immune response related proteins. Notably, differential expression changes in the proteins involved in immunity can already be observed after 10 – 12 minutes of ischemia (group *T*_2_), which shows that the alterations occur rapidly once the cells are put under ischemic conditions. These changes in immune response were mostly (20 of 34 proteins) due to a synchronous upregulation of the protein expression in both tumour and normal tissue from one time point group to the next. There is almost no differential expression effect in this group based on a tissue effect. These pathways reveal a coping mechanism involving decrease in expression of proteins that are either not survival-critical under metabolic stress induced by ischemia (such as proteins involved in drug metabolism like UGT2B17 or UGT1A8) or those that are energy intensive for the cell (ribosome biogenesis). Ischemia very interestingly appears to slowly breach through this coping mechanism by causing a decrease in tumour survival-critical proteins such as NUDT1, ELOVL, HPGDS or DZIP3 [8, 9, 10, 11].

**Table 5:** Pathways with differential protein expression *T_k_*_+1_ − *T_k_*.

| Pathways | Number of proteins |  |
| --- | --- | --- |
|  | Upregulated | Downregulated |
| Immune response | 34 | 1 |
| Metabolic processes | 7 | 7 |
| Transport of molecules | 5 | 2 |
| Regulation of cell signalling | 8 | 3 |
| Cell structure and adhesion | 6 | 3 |
| Total | 60 | 16 |

#### 2.2.2 Confounder analysis

We next analysed the influence of ischemia time on differential biomolecule expression compared to other independent variables of known clinical importance by performing a classical confounder analysis using multiple linear regression. We computed a multi-variate linear model with a Gaussian error distribution at the biomolecule level for each omic type and each of the differentially expressed biomolecule sets obtained from the initial selection step described in section 2.1.

The variation of the fold-change between tumour and normal tissue expression was modelled as dependent variable to be explainable by the independent variables (details see section 4). This analysis is only a rough approximation because at the biomolecule level both statistical assumptions for Gaussian linear models of normal error distribution and linear variable relations are not fully met as revealed by statistical testing (not shown). Nevertheless, the analysis revealed informative trends when we determined the distributions of variable effect estimates for biomolecules with any statistically significant (*p <* 0.01) variable. The corresponding plots of regression coefficient estimate means, split by positive and negative effects, for the CRC cohort are shown in Figure 3. Note that variables which did not have any significant effect on any biomolecule are not shown in the confounder plots (details see section 4).

**Figure 3:**
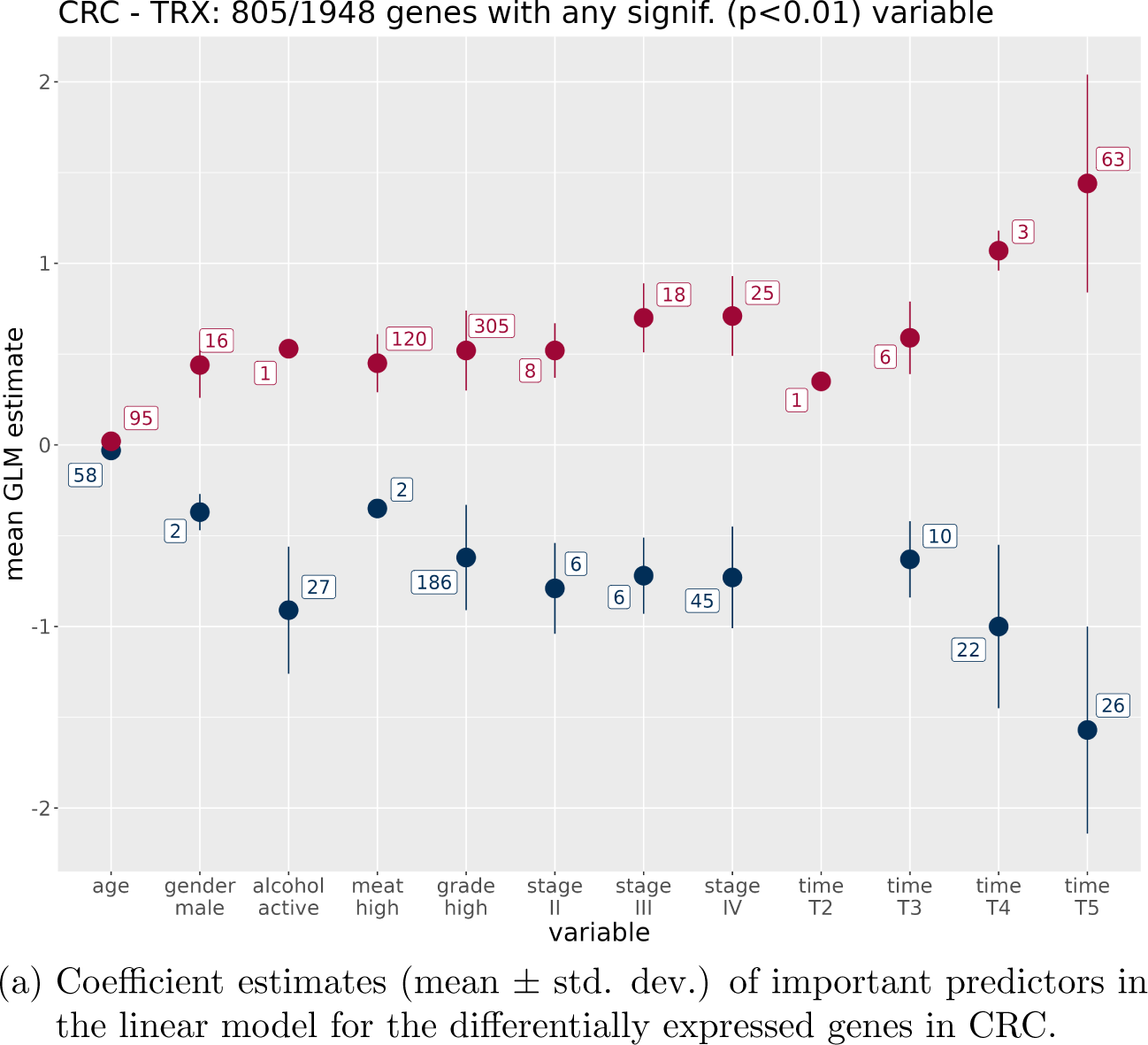
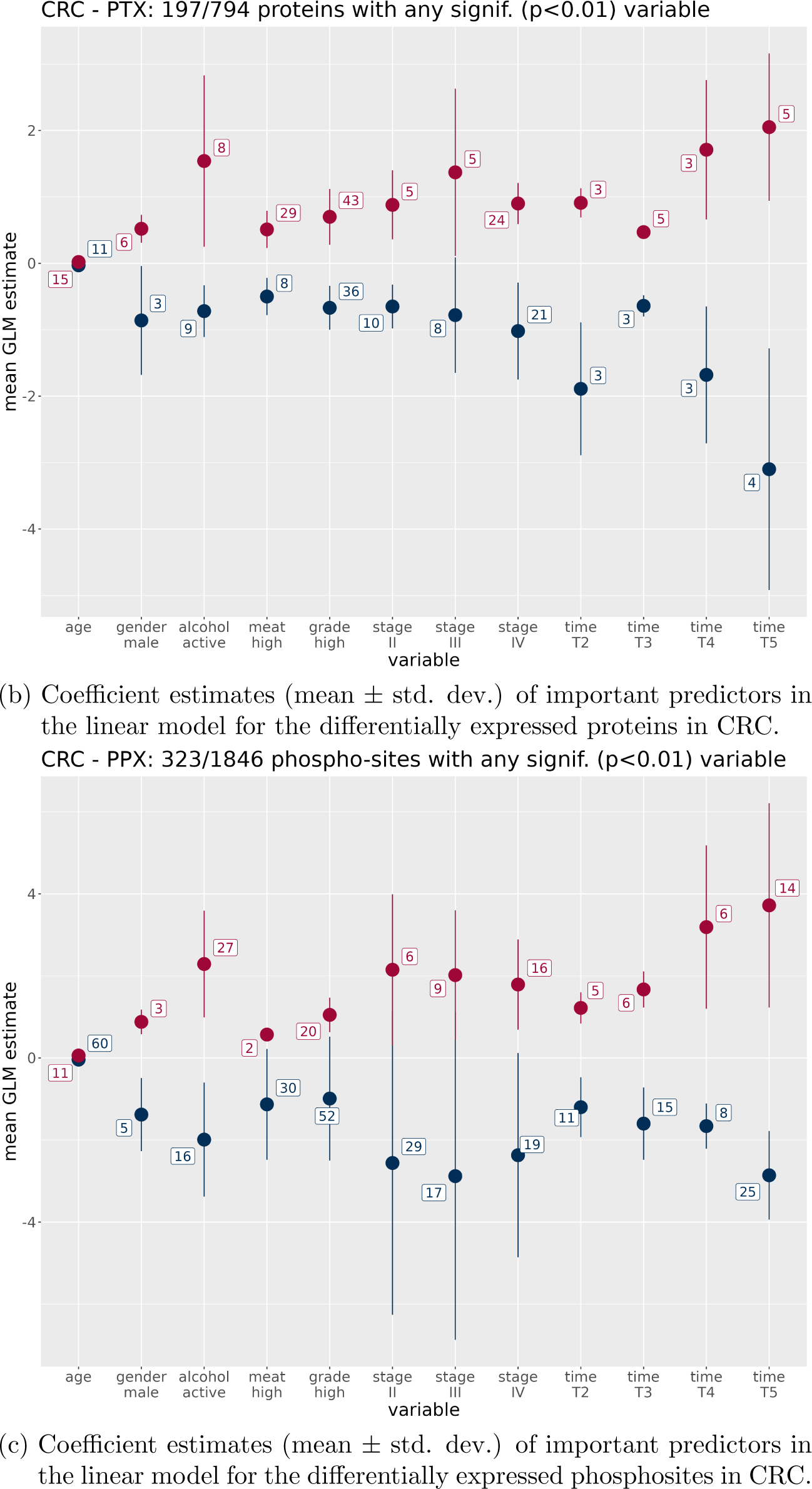
Regression coefficient estimate means and standard deviations of important predictors (linear model T-statistic with *p <* 0.01) for the differentially expressed biomolecules of the three omic modalities in CRC. Numbers related to each variable denote the number of biomolecules with any significant positive (red) or negative (blue) effect. For categorical variables, the reference variables of the linear model are indicated in section 4. For example, *T*_1_ is the reference group for the other ischemia times. (a) Transcriptome, (b) Proteome, (c) Phosphoproteome.

The figure shows that for CRC, disregarding the ischemia time variable effects (for which *T*_1_ is the reference variable), the tumour grade and stage as well as the alcohol consumption status of the patients relative to their respective references are the strongest predictors of differential biomolecule expression as expected [12, 13]. High grade has not only high coefficient estimates, but also the highest number of biomolecules with significant effects in each modality (491, 79, and 72, for mRNA, protein and phosphosite, resp.).

What about the effect of ischemia time? With the exception of protein expression, the ischemia group *T*_2_ shows the lowest *T*_1_-relative impact on the outcome compared to the other time groups, but overall, the influence of ischemia on differential biomolecule expression increases with longer ischemia times. If the ischemia time exceeds 12 minutes, the average coefficient estimate of this variable is at least as strong as the stage-IV estimates, and above 15 minutes it is much higher (also higher than grade). The number of biomolecules with significant *T*_5_ estimates also surpasses the number of biomolecules with significant stage-IV estimates, but does not exceed the number of biomolecules significantly affected by grade - which demonstrates the high relevance of this variable to explain the difference between tumour and normal tissue.

The ischemia effects gain importance from transcriptome to proteome and are strongest in phosphoproteome. But in all modalities, the magnitude of the ischemia time estimates wipes out the impact of grade and stage, which are clinically the most important predictors of cancer survival. Though this analysis is only indicative due the lack of full modelling adequacy, it provides important insights into the influence of ischemia time on biomolecule expression.

As a next step, we analysed the data to identify an ideal ischemia time cut-off.

### 2.3 Ischemia time cut-off

To find an ischemia time cut-off that optimises the trade-off of collecting the maximum amounts of samples while not impeding the identification of differentially expressed biomolecules, we investigated how many biomolecules which are differentially expressed in the short ischemia time group (< 10 min.) get lost with rising ischemia time. Figure 4 shows the relative biomolecule loss in proportion to ischemia time in the CRC cohort. As described in the methods section, we computed the number of differentially expressed biomolecules exclusive to the shortest ischemia time group as compared to the other groups for the three omic modalities, and observed overall a stronger biomolecule loss in groups of longer ischemia durations. For mRNA, the loss at 20 minutes is 22%, but is much stronger pronounced at protein (53%) and phosphoprotein (86%) level in the CRC cohort.

**Figure 4:**
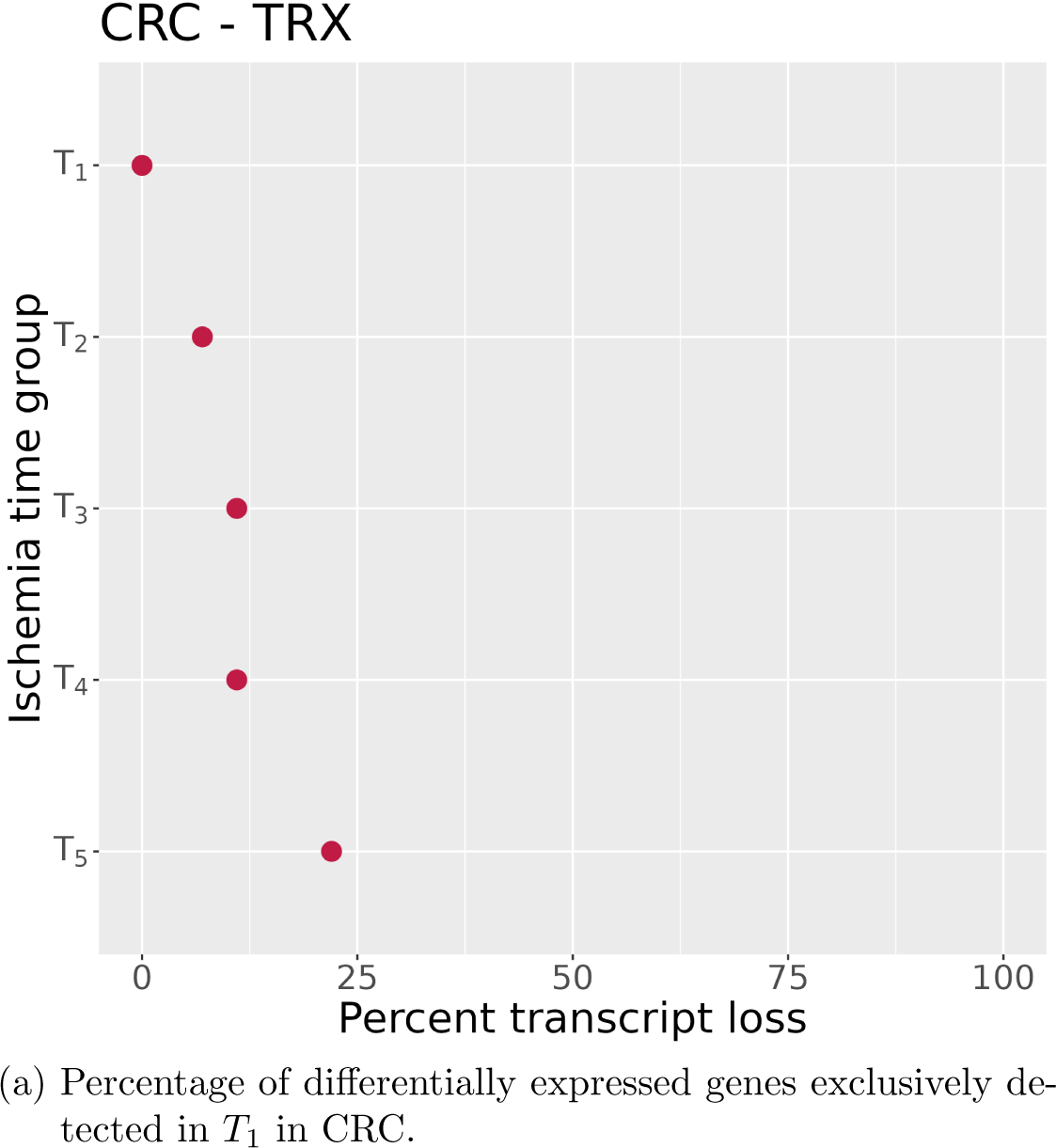
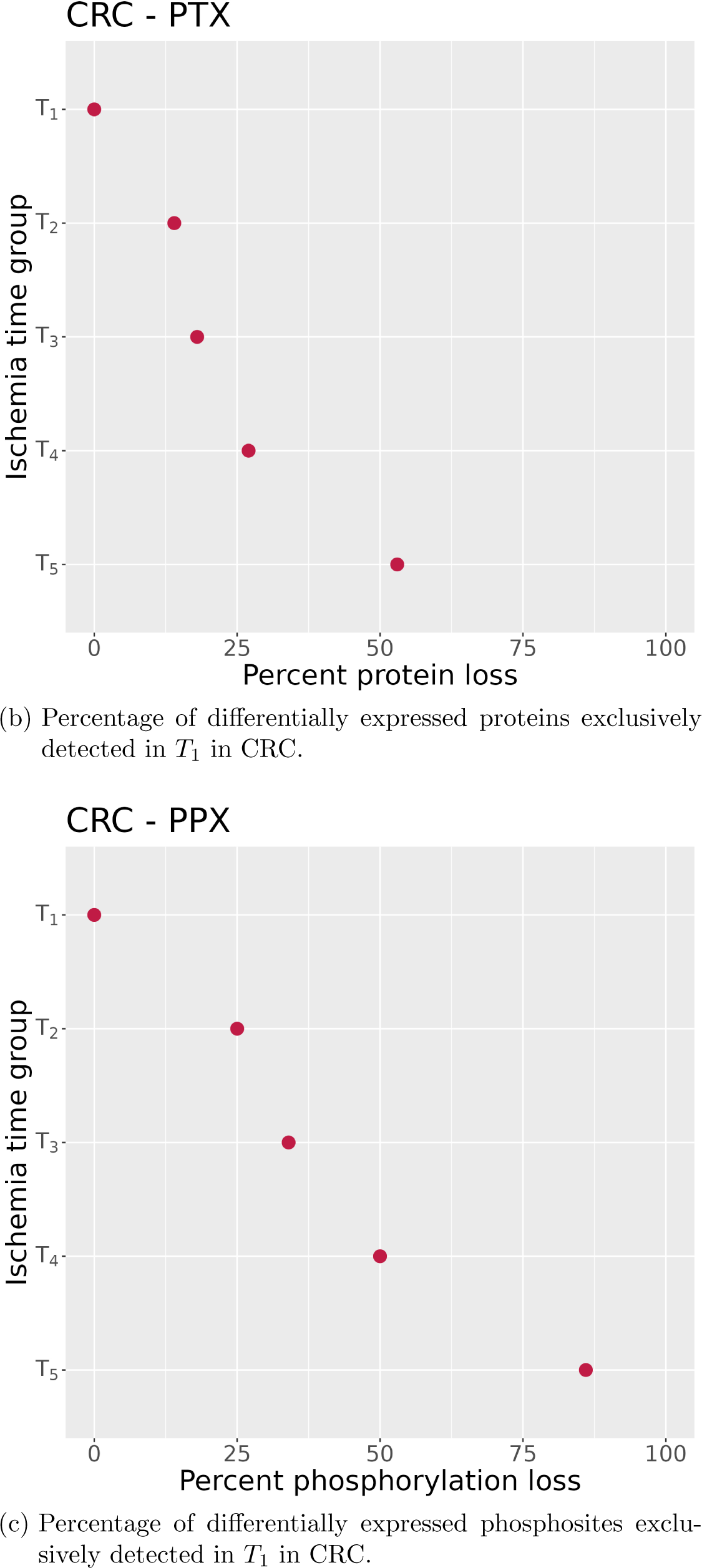
Relative biomolecule loss in proportion to ischemia times in CRC. Dot plots showing the percentage of differentially expressed biomolecules exclusively detected in the shortest ischemia time group *T*_1_ (details see section 4) for each modality. The ordinate indicates the time group compared to *T*_1_, ordered from shortest to longest time interval, the abscissa indicates the percent of biomolecule loss.

### 2.4 Results for other cancer types

We next compared the most important results from the CRC analysis to the effects of ischemia time on differential biomolecule expression in HCC, LUAD, and LUSC tumour versus matching normal tissue. Because of small group sizes (*T*_3_ and *T*_5_) in HCC (see Table 1), we adjusted the time groups we used for the other cancer types in order to enable a reasonable statistical pseudo-time series analysis. Table 6 shows the adapted classification scheme which was applied in the HCC cohort; note that due to the different assignment to the groups, there is no group *T*_5_. Figure 5 shows the confounder analysis for HCC.

**Figure 5:**
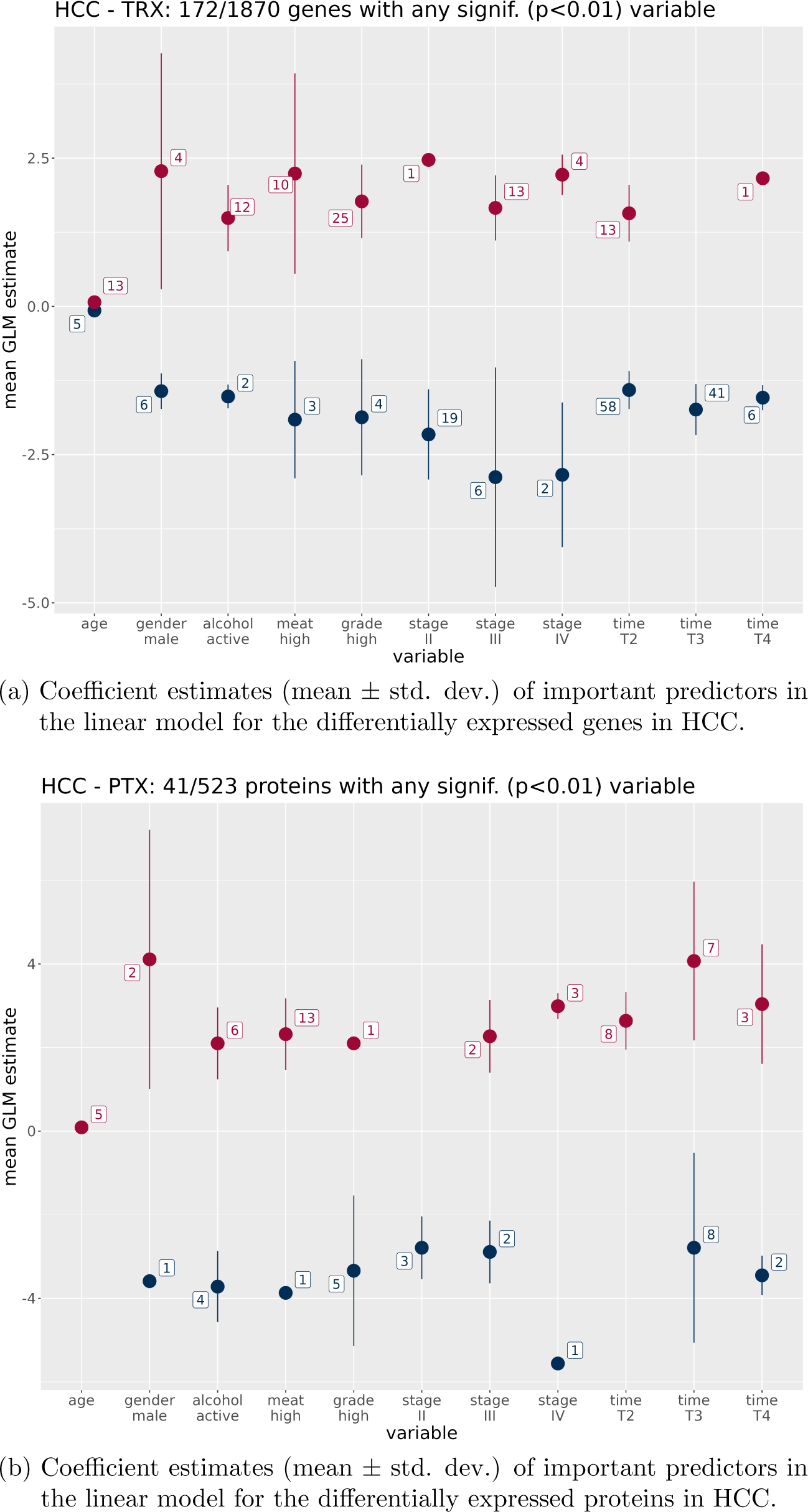
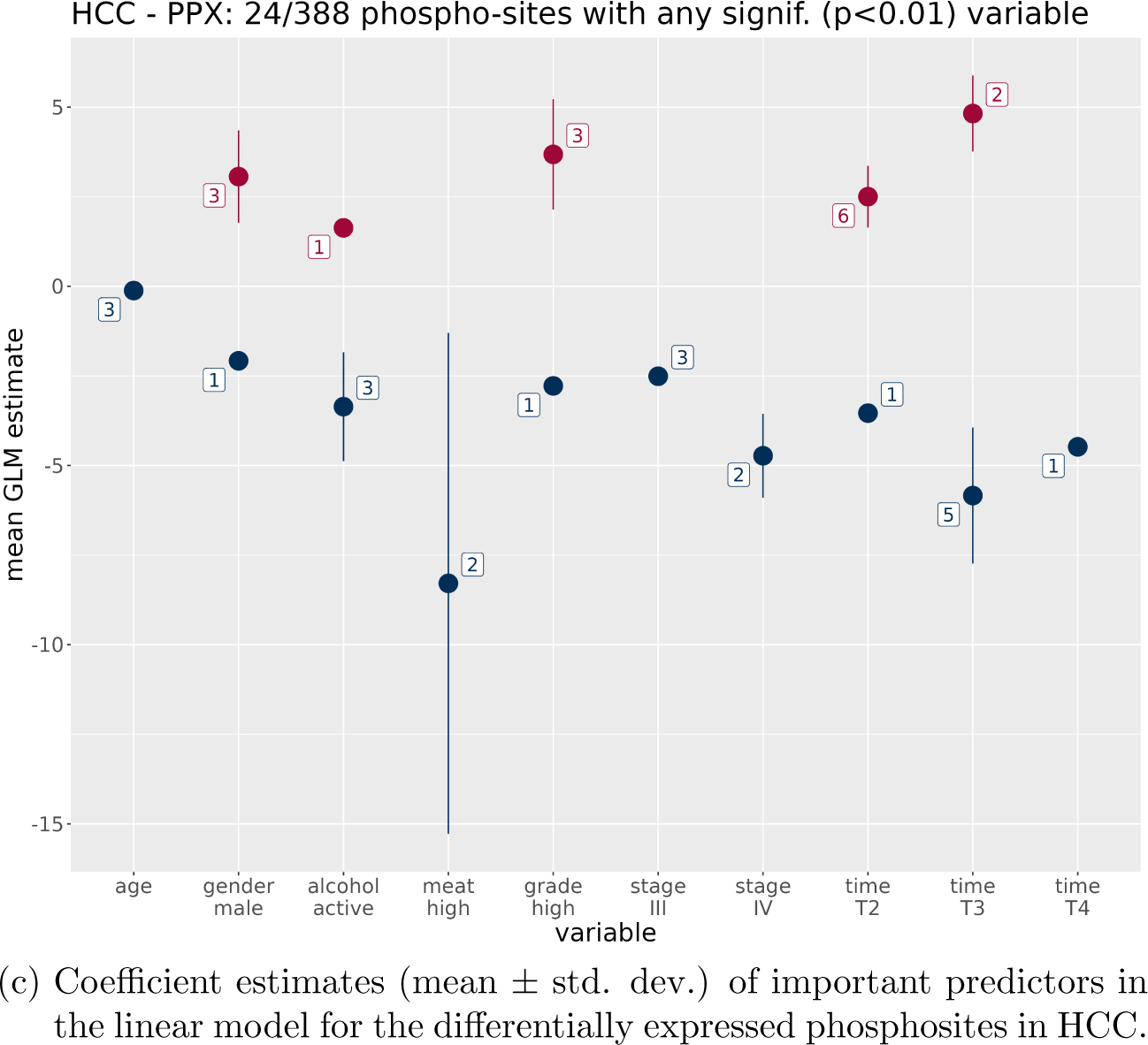
Mean coefficient estimates and standard deviation of important predictors (linear model T-statistic with *p <* 0.01) for the differentially expressed biomolecules in HCC. Numbers _1_r_7_elated to each variable denote the number of biomolecules with any significant positive (red) or negative (blue) effect. For categorical variables, the reference variables of the linear model are indicated in section 4. (a) Transcriptome, (b) Proteome, (c) Phosphoproteome.

**Table 6:**
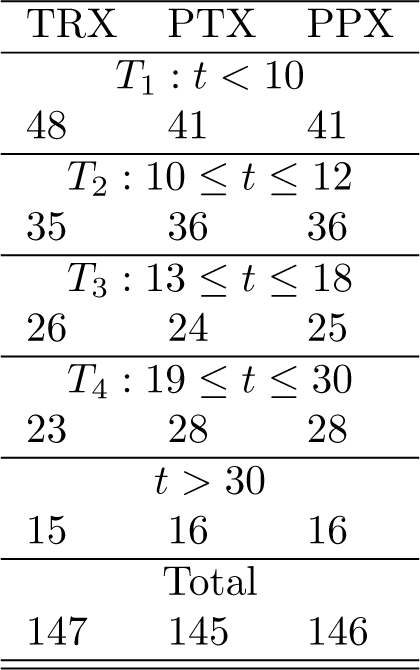
Number of tumour samples per adapted ischemia time interval (min.) with specific omic data for HCC.

Interestingly, unlike in CRC, the ischemia estimates do not surpass those of the important clinical predictors in any modality in general, though more biomolecules tend to be significantly affected by ischemia time than by stage or grade. As in CRC, the effects are more pronounced for protein and phosphoprotein modalities than for the transcriptome. In the biomolecule loss analysis for HCC (Supplementary Figure S4), the *T*_4_ group does not show a stronger loss of deregulated proteins or phosphoproteins compared to the previous time group *T*_3_. This could be explainable by the smaller group sizes of HCC samples. Nevertheless, there is an analogous trend of losing deregulated biomolecules from *T*_1_ to *T*_3_ in a monotone manner.

The confounder analysis in LUAD (Supplementary Figure S2) shows a pattern similar to CRC. Over time, the ischemia effect gains the same importance as the effect of the worst stage (mRNA) or surpasses it (protein and phophoprotein). The biomolecule loss in LUAD is comparable to CRC as well (see Supplementary Figure S5).

The confounder analysis in LUSC (Supplementary Figure S3) also shows the ischemia effect gaining importance over the clinical predictors with longer time, but in the protein and phosphoprotein modalities, there is an interesting exception, the stage IV variable. Late stage of LUSC alters differential biomolecule expression in these modalities much more than any other predictor. This is not visible in the other cancer types, though tumour grade is a very strong predictor in HCC for differential phosphosite expression. The biomolecule loss in LUSC is comparable to CRC and LUAD as well (see Supplementary Figure S6).

## 3 Discussion

There is an ongoing debate about the influence of ischemia time on data from various high-dimensional molecular characterisation modalities (‘omics’) used in the analysis of cancer tissues. This discussion is important because these data are the foundation of the identification of indicators of disease outcomes (prognosis) and novel cancer treatment targets leading to potential therapeutics.

According to some authors, ischemia times of 30 to 60 minutes can be tolerated still yielding acceptable gene expression data [2] (*n* = 6 samples). Others reported that most phosphoproteins are stable over time [3] (*n* = 3 samples) or that RNA does not degrade for two hours at room temperature [4] (*n* = 18 samples). All of these studies have an extremely limited number of samples in common which, given the high dimensionality of molecular characteristics, leads to a very low statistical power to detect ischemia time effects. On the other hand, several studies have shown that longer ischemia times reduce the quality of the measured molecular characteristics such as mRNA, protein and phosphoprotein significantly [14, 15, 16, 17, 18, 19]. However none of these studies have been able to use sufficient numbers of samples and therefore have an insufficient power to define a cut-off value for detecting the impact of ischemia time on the molecular composition of the materials, in particular for multi-omics data-driven target discovery. Therefore, so far, the published results on ischemia time have not been conclusive.

Our research demonstrates the impact of ischemia time on the relative and absolute quantities and properties of DNA, mRNA, protein and phosphoprotein molecules using a large tissue sample collection between 145 (HCC) and 633 cases (CRC) of four different cancer types in total including matching adjacent normal tissue. Because our aim is to understand the impact of ischemia time on differential biomolecule expression, we first established a set of differentially expressed biomolecules at the shortest ischemia time group *T*_1_, which most closely mimics the in vivo status. We used this baseline to map changes in these biomolecules over time.

We first consider the underlying changes of the absolute biomolecule expression in normal and tumour tissue under ischemic conditions. Table 4 shows the drastic loss in expression levels of biomolecules which are among the 5% least or most strongly expressed ones in mRNA, protein and phosphoprotein omic modalities separately for tumour and normal tissue. We must imagine this decomposition of molecules in the dying cells as highly chaotic non-ergodic complex process, during which an animate system is transformed to inanimate, decaying biomolecule matter. Fundamentally, the decay process is comparable in both tumour and normal tissues as is also evidenced by the low proportion of interaction effects (Table 3B). Therefore, the differential biomolecule expression, which compares tumour and normal tissue expression, is less affected by ischemia than the individual expression levels per tissue type.

From Figure 4 it is clear that the effect of losing differential mRNA expression over time is weaker than for proteins and phosphoproteins, though at the separate tissue levels, mRNA and protein decay in a similar manner. This could be caused by a higher regularity of the mRNA decomposition between the tissue types.

Our main findings on the influence of ischemia time on differential biomolecule expression in the CRC cohort differ between the specific omic modalities. First of all, we do not see any effect on genomic DNA, as is expected from the biochemical properties of this molecule type which can even be recovered from paleontological fossils to obtain genetic sequences.

For the other modalities, as we see for phosphoprotein (Figure 1, Table 3), but also for mRNA (Supplementary Table S1) and protein (Supplementary Table S2), ischemia times over 25 minutes make analyses of rapidly decaying molecules scientifically unattractive.

Regarding the impact of the tissue type on biomolecule expression, i.e. tumour vs. normal tissue, and the ischemia time group or the combination of both variables (interaction effect), the tissue main effect dominates the results since the biomolecules were selected based on group *T*_1_. Together with group *T*_2_ (10 ≤ *t* ≤ 12), which overall has an expression pattern very similar to *T*_1_ at least for mRNA and proteins, these time groups cover roughly ⅔ of the samples (cf. Table 1), hence dominating the main effect on biomolecule expression. The time and interaction effects in both panels of Table 3 reflect the influence of ischemia time on differential biomolecule expression. In our tissue collection, only ⅓ of the samples have longer ischemia times and their impact on expression observed at the shorter times is limited. Thus we do not see too many biomolecules with time effects due to the high quality of our tissue collection.

Our enquiry concerning shorter ischemia times shows that the interpretation of time series patterns of differential expression is challenging. The most clear pattern we observe across the time series clusters relates to immune-response proteins and metabolic function bearing proteins.

To reveal independent variables which could dominate differential expression in addition to ischemia time, we performed a systematic confounder analysis. As Figure 3 shows, increasing ischemia times even surpass the effect estimators of alcohol consumption, grade, and cancer stage in the CRC cohort, which are known to strongly affect biomolecule expression in cancer tissue [12, 13]. However, the number of biomolecules for which the grade-covariable is significanctly correlated to differential expression is not surpassed by the ischemia effect. While this can be related to the composition of our cohort (with relatively few samples with longer ischemia times, see above), the confounder analysis highlights the biological importance of grade, a predictor that is debated and sometimes underestimated in clinical practice [20].

The results obtained in the other two epithelium-descendend cancer types we analysed (LUAD and LUSC) are very similar to CRC. The number of biomolecules whose differential expression is significantly correlated to grade is also high in these cancer types.

Interestingly, in HCC, a parenchymatous cancer type, the influence of ischemia time on the differential expression outcome is weaker than in the adeno-carcinomata. However, as shown in the loss analysis (see Figure S4), there is a considerable loss of differentially expressed biomolecules in hepatic tissue was well. Taken together our findings confirm that HCC and liver normal tissue are more stable against ischemia than epithelial tissues and the cancers derived from them, so that even under longer ischemia time, there is somewhat less loss of information.

In summary, our experiments show that ischemia times below 12 minutes are recommended to obtain optimal differential biomolecule expression data. If samples with longer times are still to be included for specific reasons, they should be limited to a small proportion of the collection in order to obtain data of relevant scientific value.

## 4 Material and methods

### 4.1 Tissue sources and preparation

Indivumed GmbH has a tissue collection of fresh frozen tumour samples with matching normal tissues. These samples were frozen after different ischemia times after removal from the situs of surgery.

Tissue samples were collected by Indivumed’s clinical partners using a standardized, IRB approved protocol, focusing on minimal ischemia time. They were processed and pathologically assessed as previously described [21]. Nucleic acid extraction, library preparation and NGS were performed as previously described [21]. Protein extraction and MS analysis were performed as previously described [22, 23].

The data for CRC with matching colon mucosa from the same patient include 613 TRX (mRNA), 633 PTX (protein) and 626 PPX (phosphoprotein) data sets, for HCC with matching normal liver tissues 147 mRNA, 145 protein and 146 phosphoprotein data sets, and for LUAD 527 mRNA, 625 protein and 620 phosphoprotein data sets and for LUSC 377 mRNA, 415 protein and 408 phosphoprotein data sets with matching normal lung epithelium analyses.

Tumour stages, which are relevant for the confounder analysis we performed (see Fig. 3 and corresp. text), match the expected distribution for colon cancer surgery specimens (10% stage I, 33% II, 33% III and 20% IV). The proportions in HCC are roughly 4 : 2.5 : 2.5 : 1 for stages I to IV, and the proportions for LUAD are similar. For LUSC, they are roughly 3 : 3 : 3 : 1, inbetween CRC and HCC proportions.

### 4.2 Data processing and analysis

#### 4.2.1 Processing and normalisation

For DNA sequences obtained from whole genome sequencing, we analysed protein aminoacid sequence affecting somatic mutations (PAM, counts per locus), copy number variations (binomial distribution per genomic locus for amplification or deletion, in two matrices resp.), and the presence of truncations (binomial data). A Kruskal-Wallis test was used to identify genes with a significant mutation load on the PAM count data, Fisher’s exact test was used for this purpose on the binomial data. For each genomic locus, the resulting P-Value for each ischemia time interval was compared to the data of the shortest time interval. For mRNA sequencing data, we adjusted batch effects of sequencing providers using ComBat-seq [24]. The normalised counts were then transformed to standard TPM values (transcripts per million) accounting for the length of the transcripts as described in [25]. Protein MS/MS signals were transformed to counts using DIA-NN [26], phosphoprotein MS/MS using Spectronaut 13 (Biognosys). For both modalities, a median normalisation was performed per analysis run to scale the intensity values and make the runs comparable [27].

This preprocessing and normalisation results for each cancer type (CRC, HCC, and NSCLC) in three matrices, i.e. one matrix per omic data type (mRNA, protein and phosphoprotein), with biomolecules (genes, proteins, phosphosites) in rows and samples in columns. When needed, the expression values of the paired samples were aggregated to log_2_-fold changes (tumour versus normal tissue), yielding matrices with half the number of columns.

#### 4.2.2 Statistical analysis

*Differential biomolecule expression* was calculated for the short ischemia time group *T*_1_ using the two-sided Wilcoxon test for paired samples. P-values were adjusted for multiple testing via the method of Benjamini and Hochberg [28] which was the default correction method if not reported otherwise. The effect was computed as the mean difference of the log_2_ transformed expression values between tumour and normal tissue.

To assess differences in differential expression between tissue types (tumour versus normal) across ischemia time-groups, we used the *non-parametric Scheirer–Ray–Hare* test for 2-factorial designs with the normalised expression values as outcome.

To perform *hierarchical clustering* on the differentially expressed biomolecules in the short ischemia time group *T*_1_, for each modality the matrix of mean log_2_-fold changes (tumour versus normal tissue) was row-wise normalised (mean-centered and scaled by dividing by standard deviation). We then applied hierarchical clustering with complete linkage (using the Euclidean distance metric 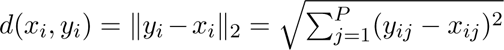, with *x_i_* and *y_i_* two matrix rows and *j* = 1 … *P* the matrix columns) to the matrix rows and displayed the results as heatmaps with dendrograms. The number of clusters was determined by visual inspection. The clusters were used to refine the time groups and limit them to 20 min.; they are merely illustrative and irrelevant for the further analyses. For the *clustering of these deregulated biomolecules in the refined ischemia time groups*, we used a Dirichlet process (DP) Gaussian process (GP) mixture model [29]. This non-parameteric time series analysis technique jointly models data clusters with a Dirichlet process and temporal dependencies with Gaussian processes.^5^ The DPGP software^6^ was parameterised with concentration parameter *α* = 0.1, the number of empty clusters in each iteartion *m* = 12, and the shape and scale parameters of the inverse gamma distribution set to *α^IG^* = 4 and *β^IG^* = 2, respectively.

The *confounder analysis* was performed by computing biomolecule-wise linear models with Gaussian error distribution for each modality using the differential biomolecule expression as response variable and *age*, *gender*, *alcohol consumption* and red *meat consumption* anamnesis, *histological grade*, *tumour stage* and the *refined ischemia time* groups as independent predictor variables. The values of the resulting effects as average changes in the log odds of the response variable associated with a one unit increase in each predictor variable were visualised, also including standard deviation, for the biomolecules with at least one significant predictor variable (t-statistic derived *p <* 0.01). Apart from the numeric continuous variable *age*, all other predictor variables were categorical ones with the following factor levels:

- *gender* : female (reference), male,
- *alcohol consumption*: inactive (reference), active,
- *meat consumption* (days per week): low (0-3 days, reference), high (4-7 days),
- *histological grade*:low (G1-G2, reference), high (G3-G4),
- *tumour stage*: I (reference), II, III, IV,
- *ischemia time*: *T*_1_ (reference), *T*_2_, *T*_3_, *T*_4_, *T*_5_.

The optimal *ischemia time cut-off* was inferred within a set-difference analysis for the refined time groups. This *set-difference analysis* was based on the results of the differential expression analysis. In detail, we identified the significantly differentially expressed biomolecules per time group *T_i_, i* = 1 … 5 (*α*_fdr_ = 0.01). We then determined the set differences *T*_1_ \ *T_i_* of deregulated biomolecules in the shortest ischemia time group and the other groups. The percentage of the number of biomolecules in the set differences in relation to the number of differentially expressed biomolecules in the shortest ischemia time group was visualised in dot plots.

## Ethics declarations

### Ethics statement

The study was approved by the local Ethics Committees, and the patients signed the appropriate informed consent.

### Author contributions

JL and SvH performed the mathematical analyses and wrote the paper. NR, AW and JLM performed the biological interpretation. Sample and clinical data collection as well as related preparation was conducted by NG, XM, TY, YT, GM and HJ.

## Footnotes

1 https://www.cancer.gov/tcga

2 Note that the differing denominator between the two groupings is due to reassigned samples and a subsequently altered pattern of missing values.

3 Note that since it is impossible to obtain patient-wise time series data based on surgical tissue removal, this is merely a pseudo-time-series.

4 https://www.genome.jp/kegg/mapper/search.html

Note that since it is impossible to obtain patient-wise time series data based on surgical tissue removal, this is merely a pseudo-time-series.

7 https://github.com/PrincetonUniversity/DP_GP_cluster

## References

1. Wellstein A. The pharmacological basis of therapeutics. Ed. by Brunton L and Knollmann B. 14th. McGraw Hill, 2023 :1335–450

2. Blackhall FH, Pintilie M, Wigle DA, Jurisica I, Liu N, Radulovich N, Johnston MR, Keshavjee S, and Tsao MS. Stability and heterogeneity of expression profiles in lung cancer specimens harvested following surgical resection. Neoplasia 2004; 6:761–7

3. Buffart TE, Oord RA van den, Berg A van den, Hilhorst R, Bastiaensen N, Pruijt HF, Brule A van den, Nooijen P, Labots M, Goeij-de Haas RR de, et al. Time dependent effect of cold ischemia on the phosphoproteome and protein kinase activity in fresh-frozen colorectal cancer tissue obtained from patients. Clinical Proteomics 2021; 18:1–12

4. Guo D, Wang A, Xie T, Zhang S, Cao D, and Sun J. Effects of ex vivo ischemia time and delayed processing on quality of specimens in tissue biobank. Molecular Medicine Reports 2020; 22:4278–88

5. Li Y, Ge X, Peng F, Li W, and Li JJ. Exaggerated false positives by popular differential expression methods when analyzing human population samples. Genome biology 2022; 23:79

6. Pak MG and Roh MS. Influence of cold ischemia time and storage period on DNA quality and biomarker research in biobanked colorectal cancer tissues. Kosin Medical Journal 2020; 35:26–37

7. Thomas P, Ebert D, Muruganujan A, Mushayahama T, Albou L, and Panther HM. Making genome-scale phylogenetics accessible to all. Protein Sci. 2022; 4218:8–22

8. Gad H, Koolmeister T, Jemth AS, Eshtad S, Jacques SA, Ström CE, Svensson LM, Schultz N, Lundbaäck T, Einarsdottir BO, et al. MTH1 inhibition eradicates cancer by preventing sanitation of the dNTP pool. Nature 2014; 508:215–21

9. Centenera MM, Scott JS, Machiels J, Nassar ZD, Miller DC, Zinonos I, Dehairs J, Burvenich IJ, Zadra G, Chetta PM, et al. ELOVL5 is a critical and targetable fatty acid elongase in prostate cancer. Cancer research 2021; 81:1704–18

10. Shao F, Mao H, Luo T, Li Q, Xu L, and Xie Y. HPGDS is a novel prognostic marker associated with lipid metabolism and aggressiveness in lung adenocarcinoma. Frontiers in Oncology 2022; 12:894485

11. Kolapalli SP, Sahu R, Chauhan NR, Jena KK, Mehto S, Das SK, Jain A, Rout M, Dash R, Swain RK, et al. RNA-binding RING E3-ligase DZIP3/hRUL138 stabilizes cyclin D1 to drive cell-cycle and cancer progression. Cancer research 2021; 81:315–31

12. Na HK and Lee JY. Molecular basis of alcohol-related gastric and colon cancer. International journal of molecular sciences 2017; 18:1116

13. Huo T, Canepa R, Sura A, Modave F, and Gong Y. Colorectal cancer stages transcriptome analysis. PLoS One 2017; 12:e0188697

14. David KA, Unger FT, Uhlig P, Juhl H, Moore HM, Compton C, Nashan B, Dörner A, Weerth A de, and Zornig C. Surgical procedures and postsurgical tissue processing significantly affect expression of genes and EGFR-pathway proteins in colorectal cancer tissue. Oncotarget 2014; 5:11017

15. Freidin MB, Bhudia N, Lim E, Nicholson AG, Cookson WO, and Moffatt MF. Impact of collection and storage of lung tumor tissue on whole genome expression profiling. The Journal of molecular diagnostics 2012; 14:140–8

16. Gajadhar AS, Johnson H, Slebos RJ, Shaddox K, Wiles K, Washington MK, Herline AJ, Levine DA, Liebler DC, and White FM. Phosphotyrosine signaling analysis in human tumors is confounded by systemic ischemia-driven artifacts and intra-specimen heterogeneity. Cancer research 2015; 75:1495–503

17. Mertins P, Yang F, Liu T, Mani D, Petyuk VA, Gillette MA, Clauser KR, Qiao JW, Gritsenko MA, Moore RJ, et al. Ischemia in tumors induces early and sustained phosphorylation changes in stress kinase pathways but does not affect global protein levels. Molecular & cellular proteomics 2014; 13:1690–704

18. Spruessel A, Steimann G, Jung M, Lee SA, Carr T, Fentz AK, Spangenberg J, Zornig C, Juhl HH, and David KA. Tissue ischemia time affects gene and protein expression patterns within minutes following surgical tumor excision. Biotechniques 2004; 36:1030–7

19. Unger FT, Lange N, Krüger J, Compton C, Moore H, Agrawal L, Juhl H, and David KA. Nanoproteomic analysis of ischemia-dependent changes in signaling protein phosphorylation in colorectal normal and cancer tissue. Journal of translational medicine 2016; 14:1–15

20. Chen K, Collins G, Wang H, and Toh JWT. Pathological features and prognostication in colorectal cancer. Current Oncology 2021; 28:5356–83

21. Yang X, Smirnov A, Buonomo OC, Mauriello A, Shi Y, Bischof J, Woodsmith J, Bove P, Rovella V, Scimeca M, Sica G1TG, Wang Y, Servadei F, Melino G, Candi E, et al. A primary luminal/HER2 negative breast cancer patient with mismatch repair deficiency. Cell Death Discovery 2023; 9:365

22. Han Y, Rovella V, Smirnov A, Buonomo OC, Mauriello A, Perretta T, Shi Y, Woodmsith J, Bischof J, Bove P, Juhl H, Scimeca M, Sica G, Tisone G, Wang Y, Giacobbi E, Materazzo M, et al. A BRCA2 germline mutation and high expression of immune checkpoints in a TNBC patient. Cell Death Discovery 2023; 9:370

23. Bruderer R, Sondermann J, Tsou CC, Barrantes-Freer A, Stadelmann C, Nesvizhskii AI, Schmidt M, Reiter L, and Gomez-Varela D. New targeted approaches for the quantification of data-independent acquisition mass spectrometry. Proteomics 2017; 17:1700021

24. Zhang Y, Parmigiani G, and Johnson WE. ComBat-seq: batch effect adjustment for RNA-seq count data. NAR Genomics and Bioinformatics 2020; 2:lqaa078

25. Wagner GP, Kin K, and Lynch VJ. Measurement of mRNA abundance using RNA-seq data: RPKM measure is inconsistent among samples. Theory in Biosciences 2012; 131:281–5

26. Demichev V, Messner CB, Vernardis SI, Lilley KS, and Ralser M. DIA-NN: neural networks and interference correction enable deep proteome coverage in high throughput. Nature methods 2020; 17:41–4

27. Kultima K, Nilsson A, Scholz B, Rossbach UL, Fälth M, and Andren PE. Development and evaluation of normalization methods for label-free relative quantification of endogenous peptides. Molecular & Cellular Proteomics 2009; 8:2285–95

28. Benjamini Y and Hochberg Y. Controlling the false discovery rate: a practical and powerful approach to multiple testing. Journal of the Royal statistical society: series B (Methodological) 1995; 57:289–300

29. McDowell IC, Manandhar D, Vockley CM, Schmid AK, Reddy TE, and Engelhardt BE. Clustering gene expression time series data using an infinite Gaussian process mixture model. PLoS computational biology 2018; 14:e1005896

