## Supplementary Figures for "Tumour specimen cold ischemia time impacts molecular cancer drug target discovery"

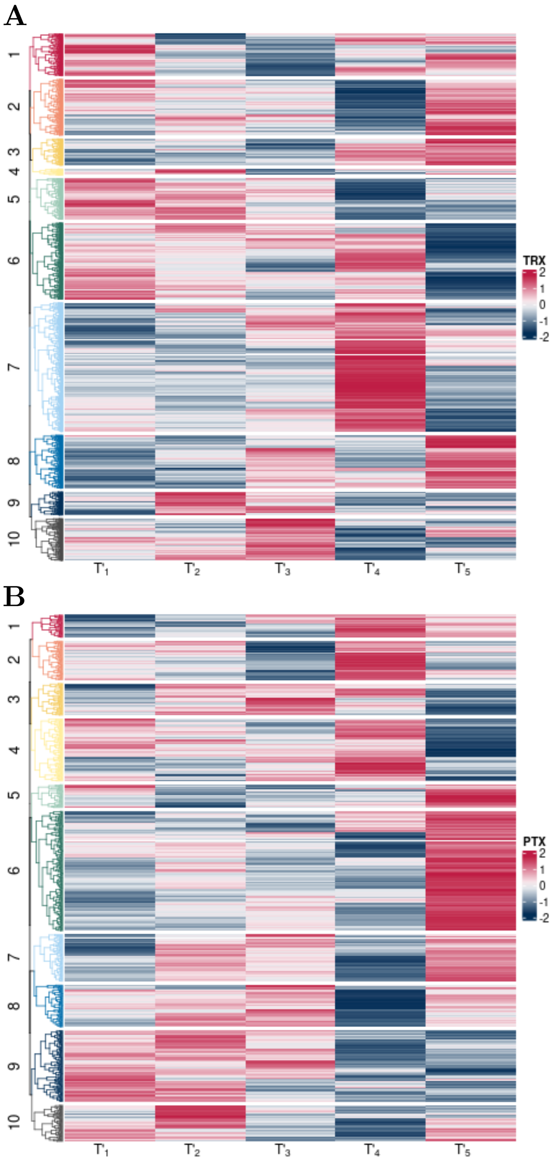


Figure S1: Differential biomolecule expression of the selected biomolecules in 5-minute intervals (abscissa) grouped into ten clusters (ordinate). Colours indicate $\log_2$-fold expression differences (biomolecule-wise standardised) between tumour and normal tissue. Red: upregulation in tumour, blue: downregulation. A: mRNA, B: Protein.

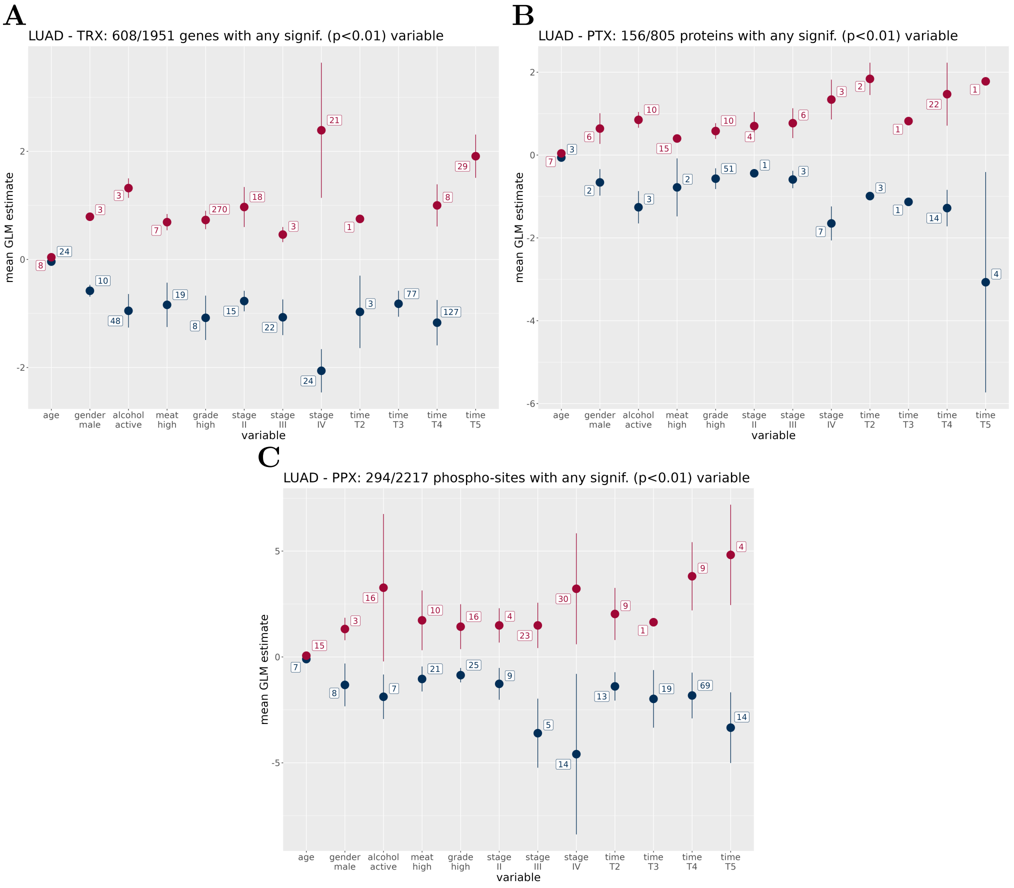


Figure S2: Distribution of mean coefficient estimates and standard deviation of important predictors (linear model T-statistic with $p < 0.01$) for the differentially expressed biomolecules of the three omic modalities in LUAD. Numbers related to each variable denote the number of biomolecules with any significant positive (red) or negative (blue) effect. For categorical variables, the reference variables of the linear model are indicated in section 4. **A:** Transcriptome, **B:** Proteome, **C:** Phosphoproteome.

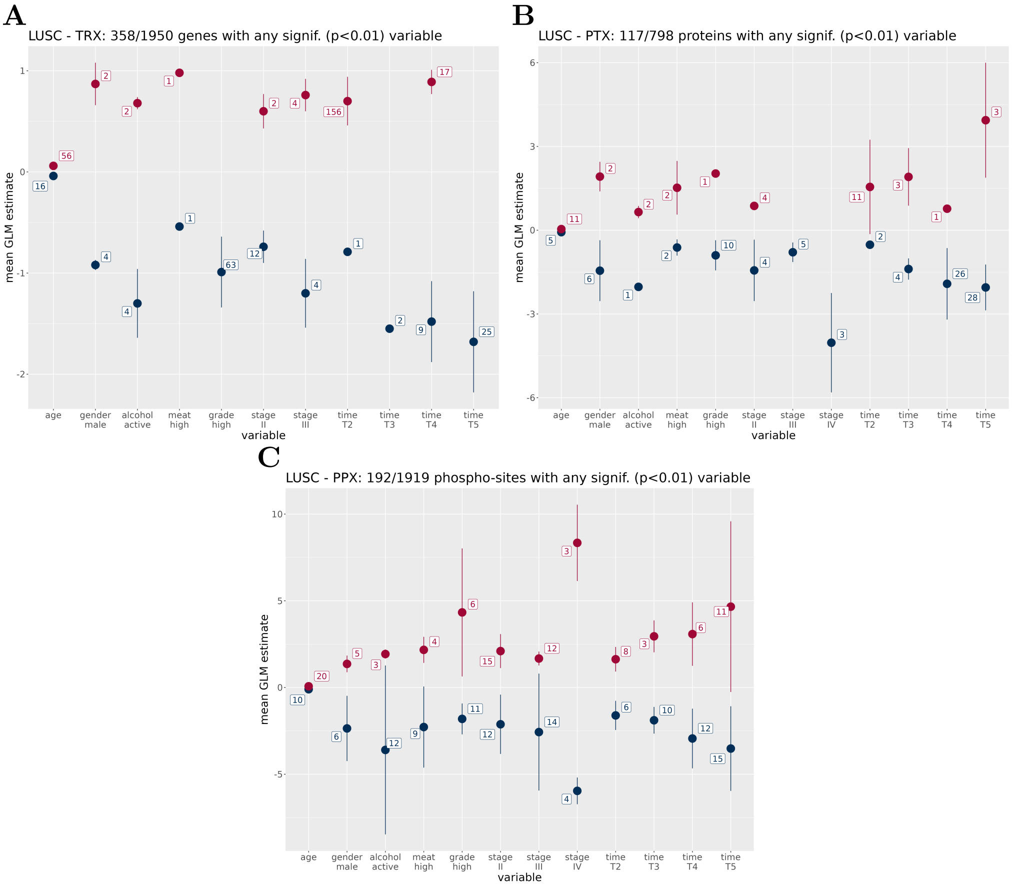


Figure S3: Distribution of mean coefficient estimates and standard deviation of important predictors (linear model T-statistic with $p < 0.01$) for the differentially expressed biomolecules of the three omic modalities in LUSC. Numbers related to each variable denote the number of biomolecules with any significant positive (red) or negative (blue) effect. For categorical variables, the reference variables of the linear model are indicated in section 4. **A:** Transcriptome, **B:** Proteome, **C:** Phosphoproteome.

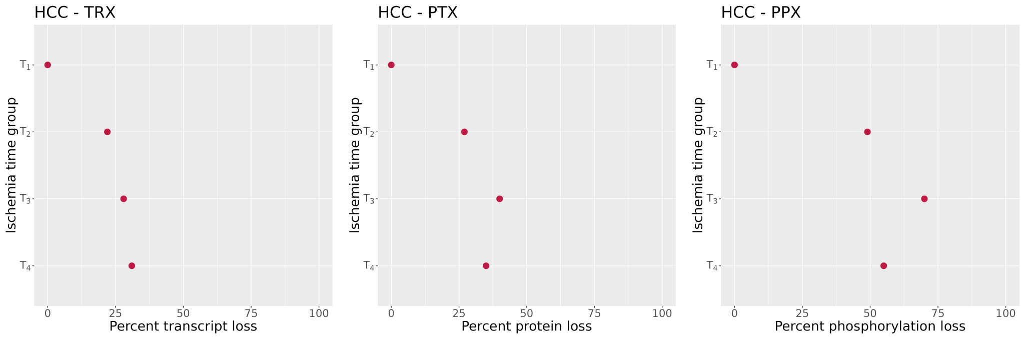


Figure S4: Relative loss of differentially expressed biomolecules in proportion to ischemia times in HCC. Dot plots showing the size of the set difference of differentially expressed biomolecules in the shortest minus all the other ischemia time groups, respectively (details see section 4) for each modality. The ordinate indicates the time group, top: shortest, bottom: longest, the abscissa the percent of biomolecule loss.

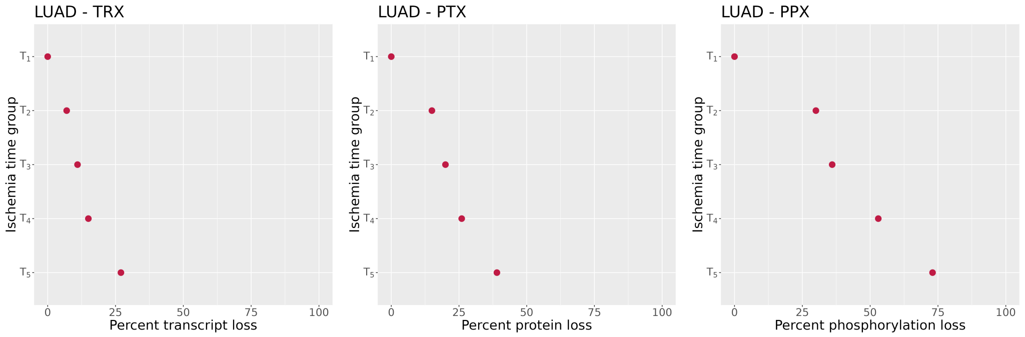


Figure S5: Relative loss of differentially expressed biomolecules in proportion to ischemia times in LUAD. Dot plots showing the size of the set difference of differentially expressed biomolecules in the shortest minus all the other ischemia time groups, respectively (details see section 4) for each modality. The ordinate indicates the time group, top: shortest, bottom: longest, the abscissa the percent of biomolecule loss.

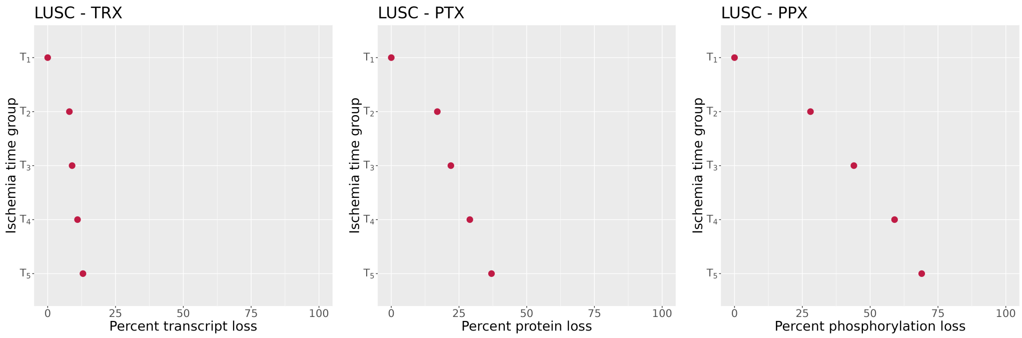


Figure S6: Relative loss of differentially expressed biomolecules in proportion to ischemia times in LUSC. Dot plots showing the size of the set difference of differentially expressed biomolecules in the shortest minus all the other ischemia time groups, respectively (details see section 4) for each modality. The ordinate indicates the time group, top: shortest, bottom: longest, the abscissa the percent of biomolecule loss.
