## Supplementary Tables for "Tumour specimen cold ischemia time impacts molecular cancer drug target discovery"

Table S1: Alterations in gene expression over time across tissue types

| **A: Long duration ischemia time groups** | | | | **B: Refined ischemia time groups** | | | |
|---|---|---|---|---|---|---|---|
| | Significant ($\alpha_{\mathsf{fdr}} = 0.05$) effect for | | | | Significant ($\alpha_{\mathsf{fdr}} = 0.05$) effect for | | |
| Cluster | Tissue | Time | Interaction | Cluster | Tissue | Time | Interaction |
| 1a | 165 | 7 | 0 | 1b | 178 | 4 | 0 |
| 2a | 220 | 10 | 0 | 2b | 67 | 0 | 0 |
| 3a | 105 | 2 | 0 | 3b | 33 | 0 | 0 |
| 4a | 25 | 0 | 0 | 4b | 305 | 6 | 0 |
| 5a | 161 | 3 | 0 | 5b | 177 | 3 | 2 |
| 6a | 300 | 4 | 0 | 6b | 44 | 1 | 0 |
| 7a | 506 | 7 | 1 | 7b | 81 | 1 | 0 |
| 8a | 209 | 6 | 1 | 8b | 136 | 2 | 0 |
| 9a | 92 | 0 | 1 | 9b | 745 | 6 | 2 |
| 10a | 165 | 4 | 0 | 10b | 182 | 3 | 0 |
| Totals | 1948 | 43 | 3 | Totals | 1948 | 26 | 4 |

Table S2: Alterations in protein expression over time across tissue types

| **A: Long duration ischemia time groups** | | | | **B: Refined ischemia time groups** | | | |
|---|---|---|---|---|---|---|---|
| | Significant ($\alpha_{\mathsf{fdr}} = 0.05$) effect for | | | | Significant ($\alpha_{\mathsf{fdr}} = 0.05$) effect for | | |
| Cluster | Tissue | Time | Interaction | Cluster | Tissue | Time | Interaction |
| 1a | 36 | 5 | 0 | 1b | 49 | 0 | 0 |
| 2a | 62 | 5 | 0 | 2b | 138 | 0 | 0 |
| 3a | 51 | 2 | 0 | 3b | 44 | 1 | 0 |
| 4a | 99 | 3 | 0 | 4b | 126 | 2 | 0 |
| 5a | 37 | 3 | 0 | 5b | 76 | 2 | 1 |
| 6a | 190 | 16 | 6 | 6b | 88 | 1 | 0 |
| 7a | 75 | 8 | 0 | 7b | 70 | 0 | 1 |
| 8a | 67 | 9 | 0 | 8b | 110 | 2 | 0 |
| 9a | 113 | 10 | 0 | 9b | 33 | 1 | 0 |
| 10a | 58 | 3 | 0 | 10b | 53 | 1 | 0 |
| Totals | 788 | 64 | 6 | Totals | 787 | 10 | 2 |
